# α-Synuclein purification significantly impacts seed amplification assay performance and consistency

**DOI:** 10.1101/2025.05.08.652548

**Authors:** Zaid A.M. Al-Azzawi, Nicholas R.G. Silver, Simeng Niu, Wen Luo, Irina Shlaifer, Martin Ingelsson, Bradley T. Hyman, Jean-François Trempe, Thomas M. Durcan, Joel C. Watts, Edward A. Fon

## Abstract

α-Synuclein seed amplification assays are a promising diagnostic tool for synucleinopathies such as Parkinson’s disease and multiple system atrophy. Standardized conditions are required to ensure a high degree of inter- and intra-laboratory reproducibility when performing these assays. A significant issue that hinders the utility of seed amplification assays is the *de novo* aggregation propensity of the α-synuclein substrate as well as inter-batch heterogeneity. While much work has focused on determining appropriate seed amplification assay buffer compositions as well as the type and amount of seed used, a robust comparison of α-synuclein substrate purification methods has not been reported. We therefore compared the utility of recombinant α-synuclein purified using four different methods as seed amplification assay substrates across two laboratories. Osmotic shock-purified α-synuclein monomer substrate showed the lowest propensity for *de novo* aggregation, which translated into being the best substrate for seed amplification assay reactions seeded with α-synuclein preformed fibrils or patient brain homogenates. Furthermore, osmotic shock α-synuclein monomer showed the best inter-batch reproducibility compared to all other substrates tested. As α-synuclein seed amplification assays continue to evolve and move towards adoption in the clinical realm, this work showcases the vital importance of standardizing the production and characterization of recombinant α-synuclein substrate. We encourage the widespread adoption of osmotic shock α-synuclein monomer as the universal substrate for seed amplification assays to maximize intra- and inter-laboratory reproducibility.

## Introduction

α-Synuclein (αSyn), encoded by the *SNCA* gene, is a presynaptic 140 amino-acid neuronal protein. While αSyn’s native function remains unclear, its aggregation is a pathological hallmark of several neurodegenerative diseases known collectively as synucleinopathies, which include Parkinson’s disease (PD), multiple system atrophy (MSA) and dementia with Lewy bodies (DLB). Seed amplification assays (SAAs) have drawn considerable attention recently as a promising diagnostic tool for synucleinopathies [1–4]. SAAs are relatively easy and cost-efficient to perform, requiring only basic laboratory technology such as a fluorescence plate reader and inexpensive, non-hazardous chemicals that are readily accessible [1]. In its most basic form, SAAs consist of recombinant IZSyn incubated under specific temperature and buffer conditions with a small amount of a seed-containing biospecimen [1]. Due to the prion-like nature of IZSyn aggregates, if the biospecimen contains an IZSyn aggregate, natively-folded recombinant IZSyn monomers will be recruited by the aggregate to undergo template-directed misfolding, resulting in propagation of the aggregate structure [1]. The kinetics of the accumulation of aggregated αSyn can be tracked in real time using the fluorescent amyloid-binding dye Thioflavin T (ThT), which selectively fluoresces when it binds to β-sheet-rich regions in protein aggregates [5].

The first publications on the use of IZSyn SAA in patient cohorts compared cerebrospinal fluid samples from PD and DLB patients to Alzheimer’s disease (AD) patients and healthy controls [6]. In these initial reports, it was demonstrated that SAAs can be useful in the diagnosis of IZ-synucleinopathy patients [6]. With further developments it has been possible to reduce the duration required to run an SAA, from durations greater than a week to only a day [7]. The specificity and sensitivity of SAAs for IZ-synucleinopathies have also been improved, with some studies reaching up to 100% and 97%, respectively [8–22]. Furthermore, it has recently been shown that SAAs may have prognostic and not solely diagnostic value [17, 22, 23].

Individual laboratories have published their own SAA protocols, with slight variations in the buffer and incubation parameters required for enabling the efficient detection and amplification of αSyn aggregates from specific biospecimens [8–22, 24–26]. As the field has progressed, the importance of standardizing assay conditions has become evident if SAAs are to be adopted for wider clinical use [27–30].

While the binary (yes/no) detection of IZSyn aggregates can be comparable between protocols, the aggregation kinetics are highly dependent on the conditions of the assay and the disease context [31]. Whereas αSyn aggregation is a shared phenomenon across synucleinopathies, their clinical manifestation, neuropathological features, and progression are distinct [32, 33]. The conformational strain hypothesis postulates that each unique synucleinopathy can be attributed to a unique structural conformation of aggregate, which is supported by current neuropathological, biochemical and structural studies [32, 33]. Distinguishing between αSyn structural conformations using SAA aggregation kinetics could facilitate the diagnosis of specific synucleinopathies [32, 33].

One group was able to distinguish PD from MSA using patient cerebrospinal fluid and brain samples via the maximal ThT fluorescence value and the lag time required for the fluorescent signal to increase. Subsequent structural and biochemical characterization confirmed these aggregated SAA end-products to be different structural polymorphs [34]. Other studies have been able to find differences in aggregation kinetics between αSyn strains, however, they often find opposite effects, which may be due to use of different SAA conditions and parameters, highlighting the lack of standardization [21, 26, 35].

While studies have investigated the effects of different buffer conditions and incubation parameters on SAA aggregation kinetics, no papers have taken into consideration the effects that changes in the recombinant IZSyn substrate can have on aggregation kinetics [21, 36]. In addition, it has been noted that variability exists between different batches of recombinant IZSyn [17, 22, 30]. However, no studies have considered the effects the recombinant αSyn purification protocol could have on the SAA kinetics, despite purification methods being a known factor that affects IZSyn aggregation propensity [36, 37].

Currently, multiple different recombinant αSyn purification protocols are used when generating the monomeric αSyn substrate for SAA [9, 11, 12, 16, 34, 38, 39]. The implicit assumption is that the properties of recombinant IZSyn produced do not vary significantly between protocols or that any effects of the purification method would be overridden by the buffer conditions used. To test these assumptions, we compared recombinant IZSyn purified by three different methods as well as a commercially sourced recombinant αSyn as substrates for SAA. The *de novo* and seeded aggregation kinetics of the four substrates were tested using SAA under standardized conditions at two different laboratory sites. We found that recombinant αSyn purified using an osmotic shock protocol serves as the most robust SAA substrate, with low rates of *de novo* protein aggregation and high inter-lab reproducibility.

## Materials and methods

### Purification of GST-tagged αSyn monomer

Glutathione S-transferase (GST)-tagged αSyn was purified as previously published [40]. A plasmid encoding GST–tagged full-length recombinant human αSyn (NM_000345) was transformed into BL21(DE3) *Escherichia coli* (New England Biolabs). When bacterial cultures reached an OD_600_ >0.6, αSyn expression was induced using 1 M isopropyl β-D-1-thiogalactopyranoside (IPTG). Induction was performed overnight after which cultures were centrifuged at 5000x g at 4 °C for 30 min. Pellets were resuspended in cold resuspension buffer (25 mM Tris-HCl pH 8, 400 mM NaCl, 5% glycerol, 0.5% Triton X-100, 5 mM PMSF, 0.5 mg/mL benzamidine, 0.5 µg/mL leupeptin, 0.5 µg/mL aprotinin, and 1 mM DTT). Resuspended pellets were then sonicated in an ice bath (5 cycles of 30 s ON/30 s OFF, 60% power). Next, the lysed pellets were centrifuged for 30 min at 18,000 rpm and 4 °C. Cleared lysates were filtered using a 0.2 µm syringe filter (VWR) and then incubated with 5 mL of glutathione Sepharose beads (GE Healthcare Life Sciences) for at least 24 h.

GST-tagged αSyn was purified using 10 mL disposable chromatography columns (Thermo Scientific) and eluted using 20 mL of freshly prepared cold elution buffer (50 mM Tris-HCl pH 8, 400 mM NaCl, 5% glycerol, 1 mM DTT, and 20 mM glutathione). Eluents were combined into an Amicon Ultra-15 centrifugal filter unit and centrifuged at 4,000x g at 4 °C for 30 min. Filtrate containing purified GST-tagged synuclein was re-suspended using 10 mL of 1X PBS and re-concentrated until the final volume of eluent was ≤ 4 mL.

GST-tagged recombinant 3C enzyme, which cleaves the GST tag from GST-tagged proteins, was expressed and purified similarly. To cleave the GST tag, GST-3C protease was added to GST-αSyn at a 1:50 mass-to-mass ratio and mixed at 4 °C overnight. Removal of the GST tag leaves 5 N-terminal linker residues on the GST-cleaved αSyn. The mixture was injected into a GSTrap 4B column (GE Healthcare Life Sciences) and the flow-through was collected into a 50 mL tube. The flow-through containing GST-cleaved αSyn was then concentrated again by centrifuging at 4,000x g at 4 °C for 30 min using an Amicon Ultra-15 centrifugal filter unit to a volume of ≤4 mL. Finally, concentrated eluent was purified using a Superdex 200 16/600 column (GE Healthcare Life Sciences) on the ÄKTA pure L system (GE Healthcare Life Sciences). The desired sample fractions were collected into an Amicon Ultra-15 Centrifugal Filter Unit, centrifuged at 3,000 rpm and 4 °C for 10 to 15 min, and adjusted to a concentration of 5 mg/mL using sterile 1X PBS. Before aliquoting into sterile tubes and freezing at -80 °C, purified αSyn was sterilized using a 0.2 μm syringe filter.

### Purification of αSyn monomer using osmotic shock

Full-length, untagged human αSyn was cloned into a pET-28a vector, expressed and purified from *E. coli* Rosetta 2 (DE3) bacteria (Novagen) using a modified osmotic shock and anion exchange protocol as follows [41–43]. When bacterial cultures reached an OD_600_ >0.6, αSyn expression was induced using IPTG for a minimum of 3 h. Next, the cells were pelleted by centrifugation at 5,000x g for 15 min at 4 °C. The cell pellet was then washed in 1X PBS and spun down using the same centrifuge parameters.

The pellet was resuspended in osmotic shock buffer (30 mM Tris–HCl pH 7.2, 40% sucrose, and 2 mM ethylenediaminetetraacetic acid (EDTA)) at 100 mL of osmotic shock buffer per 1000 mL of bacteria culture used. Following a 10 min incubation at room temperature, the suspension was centrifuged at 9,000x g for 20 min at 20 °C. The supernatant was discarded, and the pellet was quickly resuspended in ice-cold dH_2_O (40mL per 1000 mL of bacteria culture used) after which saturated MgCl_2_ (2.35 g/L) was added (40 μL per 100 mL of bacterial cell suspension) and allowed to incubate on ice for 3 min. The suspension was then centrifuged at 9,000x g for 30 min at 4°C and the supernatant was carefully collected.

Supernatant was filtered through a 0.22 μm PES filter (FroggaBio) and dialyzed into 50 mM Tris-HCl pH 8.3 using 10K MWCO SnakeSkin^TM^ Dialysis Tubing (Thermo Scientific, #68100) overnight at 4 °C. αSyn was first purified via fast protein liquid chromatography using a HiPrep Q HP 16/10 (Cytiva) anion exchange column and eluted using a linear gradient of 0 to 500 mM NaCl in 50 mM Tris-HCl pH 8.3. Fractions were assessed for purity using sodium dodecyl sulfate polyacrylamide gel electrophoresis (SDS-PAGE) and Coomassie blue staining, with fractions showing sufficiently pure αSyn being pooled and re-dialyzed into 50 mM Tris-HCl pH 8.3 overnight at 4°C. Dialyzed pooled fractions were further purified using a Mono Q anion exchange column (GE Healthcare) and eluted using the same linear gradient. Fractions were once again assessed for purity using SDS-PAGE and Coomassie blue staining. Fractions containing pure αSyn were dialyzed into dH_2_O. Concentrations of αSyn were determined by measuring absorbance at 280 nm using a NanoDrop spectrophotometer (extinction coefficient = 5,960). The protein was aliquoted into 200 μL aliquots, flash frozen in liquid nitrogen, and stored at -80 °C.

### Purification of αSyn monomer using sonication and boiling

The same bacteria, induction and harvesting procedure used in the osmotic shock protocol was replicated for the sonication and boiling protocol. Following the initial pelleting of the bacterial culture, the bacteria cell pellet was first resuspended in 10 mL of Dulbecco’s Phosphate Buffered Saline (ThermoFisher) with 50 μL of 200 mM PMSF. The cell suspension was lysed using a tip sonicator (18% amplitude, 6 pulses of 25 sec each with 2 min rest on ice in between pulses) and subsequently immersed in boiling water for 15 min. The bacterial lysate was centrifuged at 10,000x *g* for 10 min at 4 °C and the supernatant was carefully collected. The supernatant was filtered through a 0.22 μm PES filter (FroggaBio) and purified with the same two-step anion exchange procedure used in the osmotic shock protocol.

### Commercial His-tagged αSyn monomer

Recombinant full-length human αSyn monomers containing an N-terminal poly-histidine tag were purchased from Impact Biologicals (catalog #301-01).

### Circular dichroism spectroscopy

The four different preparations of recombinant IZSyn were normalized to 0.1 mg/mL and dialyzed into 10 mM phosphate buffer pH 7.4 containing 100 mM (NH_4_)_2_SO_4_. Following dialysis, samples were filtered through a 0.22 μm PES filter (FroggaBio). Circular dichroism measurements were conducted at 22 °C using a JASCO J-715 spectropolarimeter. Spectra were normalized to a blank and smoothed.

### Generation of αSyn preformed fibrils

Two separate batches of αSyn preformed fibrils (PFFs) were generated by incubating 500 μL of 5 mg/mL purified αSyn monomer (obtained using the GST-tagged approach) at 37 °C in a digital heating shaking dry bath or a thermomixer set at 1000 rpm for 5 d. At the end of the 5 days, the PFFs were sonicated for 10 cycles, 30/30 s ON/OFF at 10 °C in a water bath sonicator (Bioruptor plus, Hologic Diagenode). The preparations were then aliquoted to the required volumes (5 -100 μL) and stored at -80 °C. PFFs were further characterized for quality control using transmission electron microscopy (TEM) and dynamic light scattering (DLS) (Supplemental methods, Supplemental Fig. 1A, B).

### αSyn seed amplification assays: unseeded and PFF-seeded

0.1 mg/mL of recombinant αSyn was incubated at 37 °C in 100 µL reaction buffer (250 mM sodium citrate tribasic dihydrate pH 8.5, 80 µM SDS, 10 µM ThT in ddH_2_O). All components in the reaction mix including the recombinant αSyn were filtered using 0.2 µm syringe filters (Basix). Reactions were performed in quadruplicates in black/clear bottom 96-well plates (Corning), and underwent cycles of 29 min rest, 1 min of double orbital shaking at 400 rpm for a total of 100 h. Fluorescence readings were performed every 30 min using an excitation of 440C±C10Cnm and an emission of 480C±C10Cnm in a Fluostar Omega or a CLARIOstar microplate reader (BMG Labtech). Gain was manually set at 1,500 and orbital scanning mode was used at 3 mm. Sodium citrate buffer was compared to multiple other buffers that are commonly used and tested in unseeded vs seeded conditions including: citrate buffer at higher and lower concentrations (500 mM, 250 mM, and 125 mM), PIPES buffer (100 mM PIPES pH 6.5, 300 mM sodium chloride, 80 µM SDS, 10 µM ThT in ddH_2_O), PBS buffer (80 µM SDS, 10 µM ThT in PBS pH 7.4), and NaCl at different concentrations (300/150/75 mM NaCl, 80 µM SDS, and 10 µM ThT in ddH_2_O).

### Brain homogenate preparation

Brain homogenates (BHs) from frozen human MSA and C57BL/6 mouse tissue [10% (w/v) prepared in calcium- and magnesium-free PBS] were generated using a Minilys homogenizer and CK14 soft tissue homogenizing tubes (Bertin Corp.). Homogenates were aliquoted and then stored at -80 °C.

### αSyn seed amplification assays: brain homogenate-seeded

To generate brain-derived seeds, 10% (w/v) BH (in PBS) were centrifuged at 10,000x g for 10 min at 4 °C. The supernatant was collected, and the protein concentration in the PBS-soluble fraction was determined using the BCA assay. Reactions were performed in black 96-well plates with a clear bottom (Corning). For the reaction mixture, each well contained 10 μL of the seed (5 μg of total protein from the PBS-soluble fraction, diluted in PBS), 20 μL of 50 μM ThT (final concentration of 10 μM), 20 μL of 0.5 mg/mL monomeric recombinant αSyn (final concentration of 0.1 mg/mL), and 50 μL of the reaction buffer consisting of 80 mM phosphate buffer pH 8, 350 mM Na_3_Citrate (final concentration of 40 mM phosphate buffer pH 8, 175 mM Na_3_Citrate), and three 0.5 mm silica beads. Plates were sealed and incubated at 42 °C in a CLARIOstar microplate reader (BMG Labtech) with cycles of 1 min shaking (400 rpm double orbital) and 1 min rest for a period of 72 h. ThT fluorescence measurements (450 ± 10 nm excitation and 480 ± 10 nm emission, bottom read) were taken every 2 min. Four technical replicates were analyzed per sample.

### Analysis of seed amplification assay data

The unseeded and seeded kinetic curves were fit to a sigmoidal dose–response (variable slope) model in GraphPad Prism to obtain values for the Hill slope (k) and the time at which fluorescence is halfway between the baseline and plateau values (T50). Lag phases were then calculated using the equation T50 – [1/(2*k)] [44]. For samples that did not aggregate within the 100 h timeframe, they were assigned a lag time of 100 h. For samples that aggregated immediately, they were given a lag phase of 0 h. The *ThT_Max_* values are the highest single value reported for each individual well. Coefficient of variation (CV) was calculated by the formula CV ^a^, where σ is the standard deviation of the lag time for an independent quadruplicate and μ is the mean of the lag time for the same independent quadruplicate.

### Intact protein mass spectrometry (LC-MS)

As previously described [45], protein samples were diluted to 0.1 mg/mL in 0.1% (v/v) formic acid before 1 µg of each was injected onto a Dionex Ultimate 3000 UHPLC system at 200 µL/min using a Waters BioResolve RP mAb Polyphenyl column (450Å, 2.7 µM, 2.1 x 100 mm). The resulting eluate (5 min wash with 4% (v/v) acetonitrile in 0.1% (v/v) formic acid, followed by 20 min gradient to 90% (v/v) acetonitrile in 0.1% (v/v) formic acid) was analyzed on an Impact II QTOF mass spectrometer (Bruker Daltonics) equipped with an Apollo II ion funnel electrospray ionization source and Bruker OtofControl v4.0 / DataAnalysis v4.3 software. Following calibration (ESI-L Low Concentration Tuning Mix; Agilent Technologies #G1969-85000), data were acquired in positive-ion profile mode using a capillary voltage of 4,500 V and dry nitrogen heated at 200°C. Total ion chromatograms were used to determine where the protein eluted, and spectra were summed over the entire elution peak. Multiply charged ion species were deconvoluted at 10,000 resolution using the maximum entropy method.

### Analytical ultracentrifugation

The polymeric state of αSyn generated by multiple preparation protocols was analyzed by sedimentation velocity analysis. Samples were diluted to a concentration of 1mg/mL in water (PBS for GST-tagged IZSyn monomer) and then loaded into standard 12mm 2-sector Epon-charcoal centrifuge cells. AUC analysis was performed at 10°C in a Beckman Optima AUC analytical ultracentrifuge. Sedimentation behavior resulting from a rotor speed of 50,000 rpm (An-60 Ti rotor) was observed using the optical absorbance of each sample (A_280nm_). For each sample, two hundred radial scans were obtained at 3-minute intervals. Following calculation of temperature-corrected sample and fluid parameters using the SEDNTERP software package [46], datasets were fitted according to the continuous c(S) Lamm equation model in the SEDFIT software package (version 9.4) [47] to obtain the concentration distribution by sedimentation rate.

### Statistical analysis

All statistical analyses were performed using GraphPad Prism (v.9.3) with a significance threshold of *p* = 0.05. Data comparisons were made using either one-way ANOVA with Tukey’s multiple comparisons test, two-way ANOVA with Tukey’s multiple comparisons test, two-way ANOVA test with Šidák multiple comparisons test, a Kruskal–Wallis test followed by Dunn’s multiple comparisons test, Mann-Whitney tests with a Holm-Šidák correction for multiple comparisons, or unpaired t-tests. Under conditions where the values were restricted to an upper limit (Lag Time), non-parametric tests were used. The Mann-Whitney test was used when comparing two groups in which the assumption of non-equal standard deviations as variability between experiments was substantial, while the Kruskal-Wallis test followed by Dunn’s multiple comparisons test was used to compare multiple groups. In cases where the distribution was not restricted, unpaired two-tailed t-tests were used to compare two groups while ANOVAs were used to compare multiple groups. If post-hoc comparisons were conducted between all groups, Tukey’s multiple comparison tests were used, whereas if the number of comparisons were restricted, a Šidák multiple comparison test was used. A semilog non-linear regression was used to analyze the relationship between lag time and PFF seeding amount. Values are reported as mean ± SEM.

## Results

We compared four different preparations of recombinant full-length human IZSyn for their viability as an SAA substrate. Three of the αSyn preparations were produced “in house”, which permits greater flexibility and scalability. Recombinant αSyn monomers were purified either by fusion to a GST tag followed by purification using glutathione beads and removal of the GST tag (“<u>G</u>S<u>T</u>-tagged IZSyn <u>m</u>onomer”: GTM); by bacterial lysis via sonication and boiling followed by anion exchange chromatography (“<u>s</u>onication and <u>b</u>oiling αSyn <u>m</u>onomer”: SBM); or by bacterial lysis via osmotic shock followed by anion exchange chromatography (“<u>o</u>smotic <u>s</u>hock αSyn <u>m</u>onomer”: OSM). The fourth preparation was obtained from a commercial source and contained an N-terminal poly-histidine tag (“<u>c</u>ommercial <u>H</u>is-tagged αSyn <u>m</u>onomer”: CHM).

### Osmotic shock-purified αSyn monomer exhibits limited *de novo* aggregation

The spontaneous aggregation propensity of the four different preparations of IZSyn was tested via unseeded SAA reactions. Separate batches of the four substrates were tested at two different laboratories to ensure inter-laboratory reliability of the findings and assess the variability of the substrates (Fig. 1A, B). The GTM substrate showed the fastest *de novo* aggregation rate, with an average lag phase of 45.64 ± 6.19 h (Fig. 1C). The SBM substrate had a lag time of 74.15 ± 7.00 h, while the CHM substrate had a lag time of 86.16 ± 4.13 h, both of which were significantly longer than the GTM substrate (*p*<0.05) (Fig. 1C). Notably, the OSM substrate had the longest lag phase of 98.56 ± 1.08 h, which was significantly longer than for the other three substrates (*p*<0.05) (Fig. 1C). *ThT_Max_* values may correlate with either the structure of the aggregates formed at the end of the reaction or the extent of aggregation. The *ThT_Max_* value of the OSM was 1,922 ± 1,138, which was significantly lower than the other three substrates (*p*<0.05), congruent with the lack of *de novo* aggregation seen with this substrate (Fig. 1C). The SBM (25,260 ± 6,114) and CHM (23,696 ± 5,609) *ThT_Max_* values were significantly lower (*p*<0.05) than the GTM substrate (55,487 ± 5,190) (Fig. 1C). Only two of the 16 OSM wells showed any aggregation, whereas 11/16 SBM, 15/16 CHM and all 16 GTM wells showed aggregation (defined as a lag time less than 100 h). All five of the SBM wells which did not aggregate were reported at a single lab, suggesting interlab variability. For the two OSM wells in which αSyn aggregation was apparent, the aggregation appears to be the most aggregation-prone substrate, while OSM appears to be the least aggregation-prone substrate. CHM and SBM exhibit intermediate spontaneous aggregation kinetics. To further probe the utility of the OSM substrate, we tested multiple additional batches purified at different times in unseeded reactions and saw similar resistance to *de novo* aggregation across all batches (Fig. 1D). This indicates that low *de novo* aggregation potential is a robust, consistent property of OSM.

**Figure 1:**
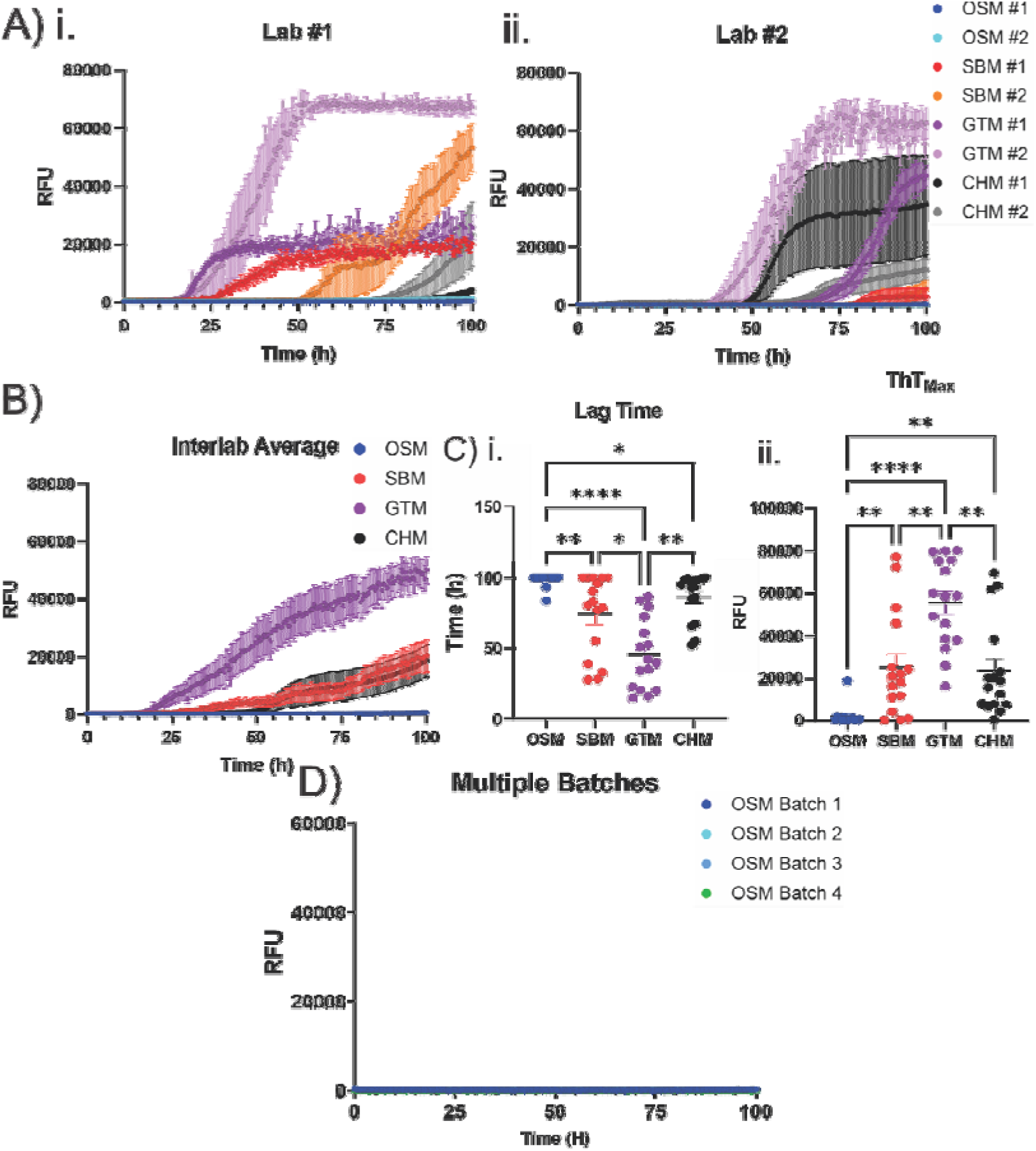
Spontaneous Aggregation Kinetics of Different Monomeric αSyn SAA. Shock Monomer (OSM), Sonicated and Boiled Monomer (SBM), GST-Tagged Monomer (GTM), or Commercially Available His-Tagged Monomer (CHM). Samples were run in quadruplicates twice in each lab (i and ii) to ensure inter-laboratory replicability of the results (n = 8 replicates per lab). Data is mean ± SEM. B) ThT aggregation curves for unseeded SAA reactions between the four trials (two from each lab), averaged to generate inter-laboratory results (n = 16 total replicates). Data is mean ± SEM. C) Comparison of the lag phase (i) and ThTMax (ii) of all the unseeded reactions (n = 16). P values for lag time were calculated using a Kruskal-Wallis test followed by Dunn’s multiple comparisons test and ThTMax values were compared using Brown-Forsythe ANOVA test with Dunnett’s T3 multiple comparisons test. Data is mean ± SEM. D) Four independent preparations of OSM were run in unseeded SAA reactions to characterize the de novo aggregation propensity of multiple batches of OSM. n = 4 replicates per batch. Data is mean ± SEM.

### Osmotic Shock Monomer Showed Dose Dependent Seeding in PFF SAA

The lower *de novo* aggregation potential of OSM compared to other recombinant αSyn preparations prompted us to investigate at one laboratory if it could be used for the sensitive and specific detection of αSyn PFF seeds in SAA. The PFFs were generated as described above using GTM and formation was verified with TEM and DLS (Supplemental Fig. 1). The OSM substrate produced positive seeding reactions when seeded with up to 10^-6^ dilutions of PFFs (Fig. 2A). There was a direct relationship between the dose of PFF added and the resultant lag time, with higher PFF dilutions resulting in significantly longer lag times in SAA, whereas the *ThT_Max_* was unaltered (Fig. 2B-D).

**Figure 2:**
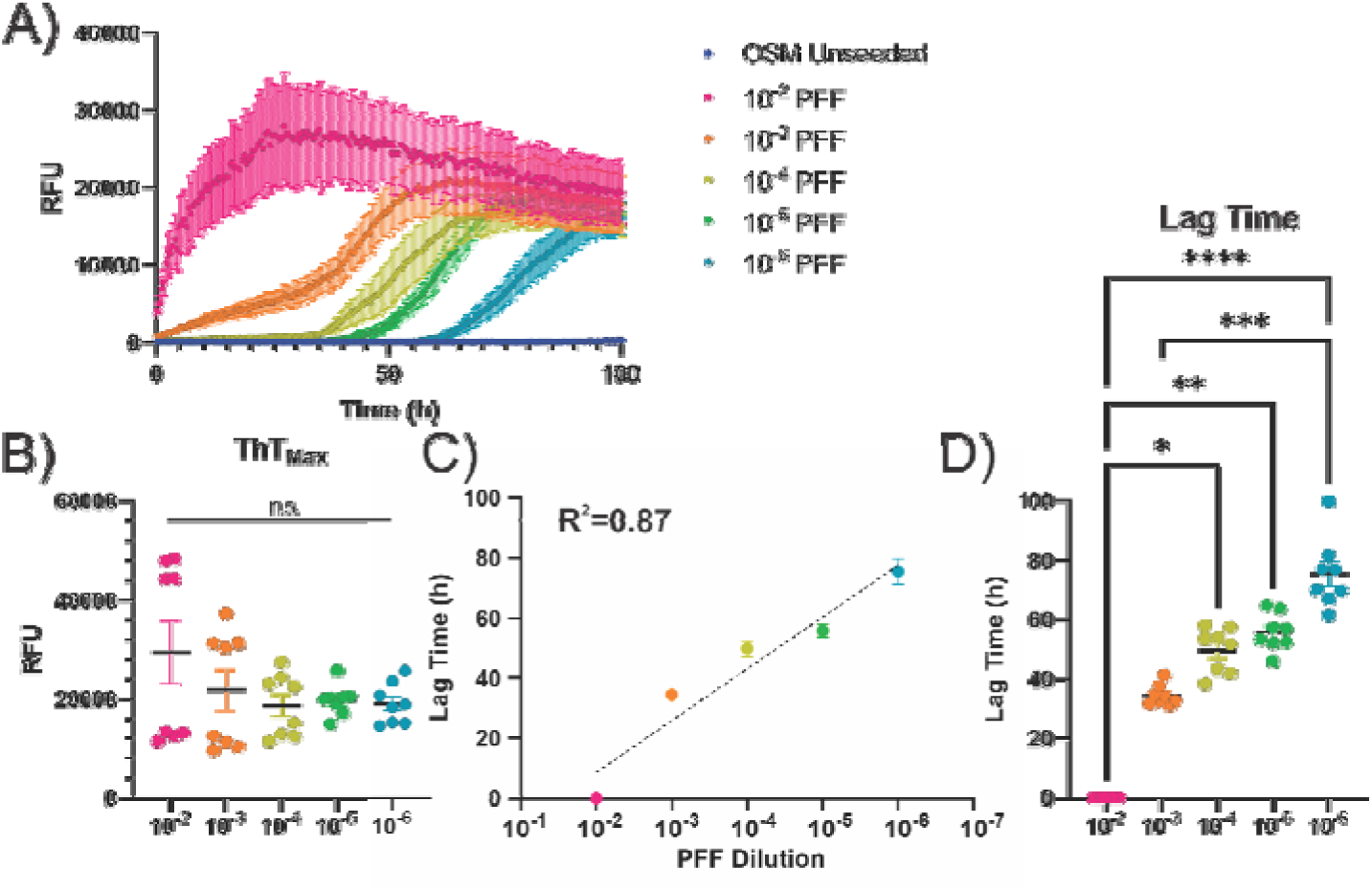
Dose-Dependent Detection of PFF Seeds Using SAA with OSM Substrate. **A)** PFF-seeded reactions show a dose-dependent delay in the kinetics of αSyn aggregation with increasing PFF dilutions. The experiment was performed twice with four technical replicates per experiment. **B)** No significant differences in final *ThT_Max_* value were observed across different PFF seed dilutions. **C)** A direct relationship between PFF seed dilutions and lag time was noted, with an R^2^ of 0.87. **D)** Lag times significantly increase with decreasing PFF seed amount. *P* values for lag time were calculated using a Kruskal-Wallis test followed by Dunn’s multiple comparisons test and *ThT_Max_* were calculated using Brown-Forsythe ANOVA test with Dunnett’s T3 Multiple Comparisons test.

### All Monomers Show Enhanced Aggregation Kinetics in PFF SAA

To compare the performance for each of the four recombinant αSyn substrates in PFF-seeded SAA reactions, 10^-3^ dilutions of PFFs (from the same batch) were chosen to seed the reactions and tested at both laboratories. As with the unseeded reactions, the same kinetic parameters were assessed. The OSM substrate had a significantly (*p*<0.05) longer lag time (38.77 ± 1.59 h) compared to all other substrates (SBM: 22.97 ± 5.212 h; GTM: 15.94 ± 4.19 h; CHM: 20.35 ± 4.34 h) (Fig. 3A-C). The *ThT_Max_* was not significantly different between any of the four substrates (Fig. 3C).

**Figure 3:**
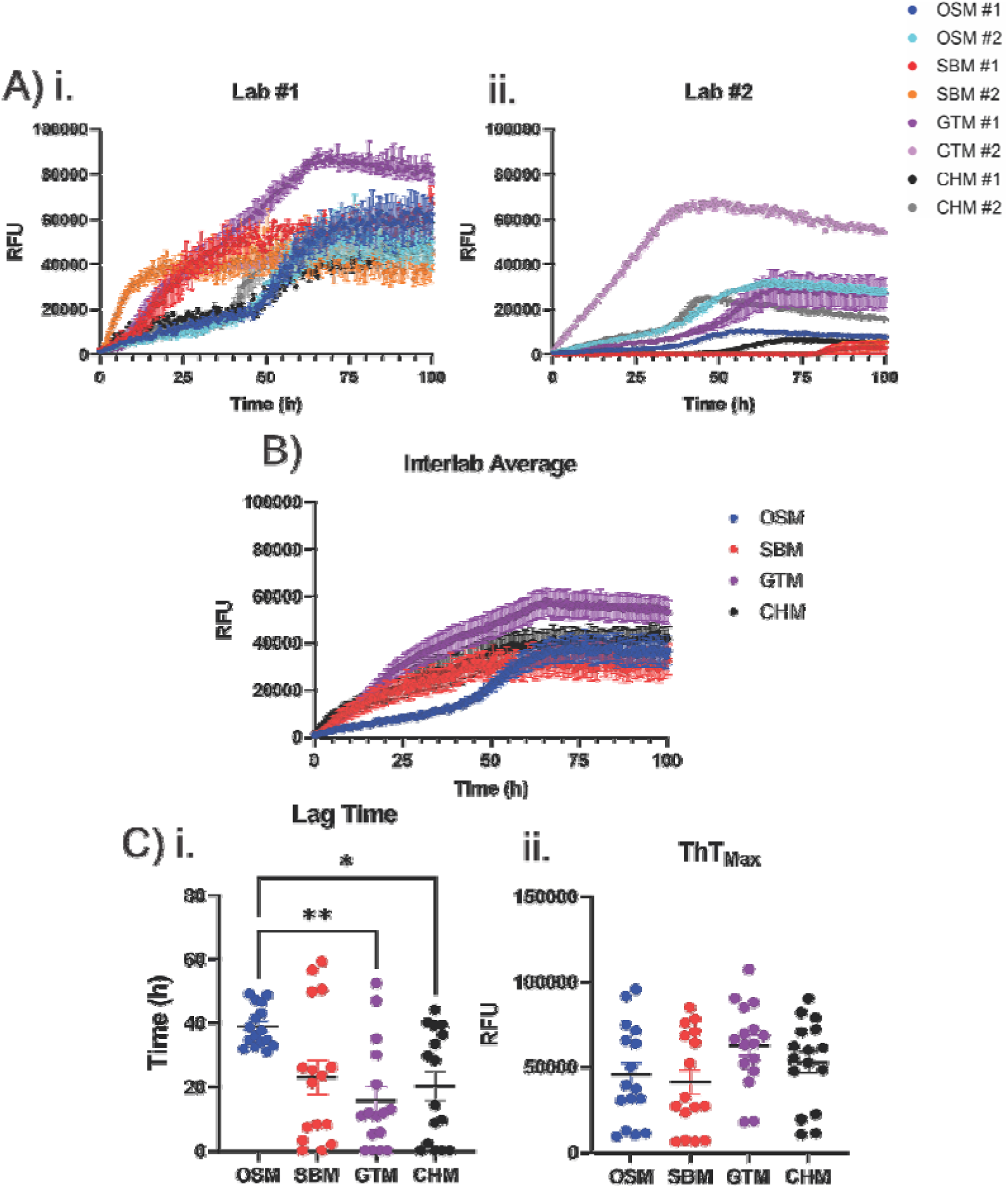
αSyn Substrate Performance in PFF-Seeded SAA Reactions. **A)** Osmotic Shock Monomer (OSM), Sonicated and Boiled Monomer (SBM), GST-Tagged Monomer (GTM), or Commercially Available His-Tagged Monomer (CHM) were seeded with 10^-3^ dilution of PFFs and run in quadruplicate twice in each lab (i and ii) to ensure the inter-laboratory replicability of the results. **B)** Results of the seeded reactions between the four trials (two from each lab) were averaged to generate inter-laboratory results. **C)** Comparison of the lag phase (i) and *ThT_Max_* (ii) of all the seeded reactions. *P* values for lag time were calculated using a Kruskal-Wallis test followed by Dunn’s multiple comparisons test and ThT_Max_ were calculated using Brown-Forsythe ANOVA test with Dunnett’s T3 Multiple Comparisons test.

When comparing the lag time of the unseeded reactions to the seeded reactions, the GTM substrate had the smallest difference between unseeded and seeded reactions of 29.71 h, while the OSM (59.79 h), SBM (51.18 h) and CHM (65.80 h) all had larger differences between unseeded and seeded reactions (Fig. 4A, B). These differences were all significant (*p*<0.05) (Fig. 4B). However, the *ThT_Max_* was only significantly different between unseeded and seeded reactions with the OSM and CHM substrates, suggesting that the end-products for the SBM and GTM may be comparable (Fig. 4B).

**Figure 4:**
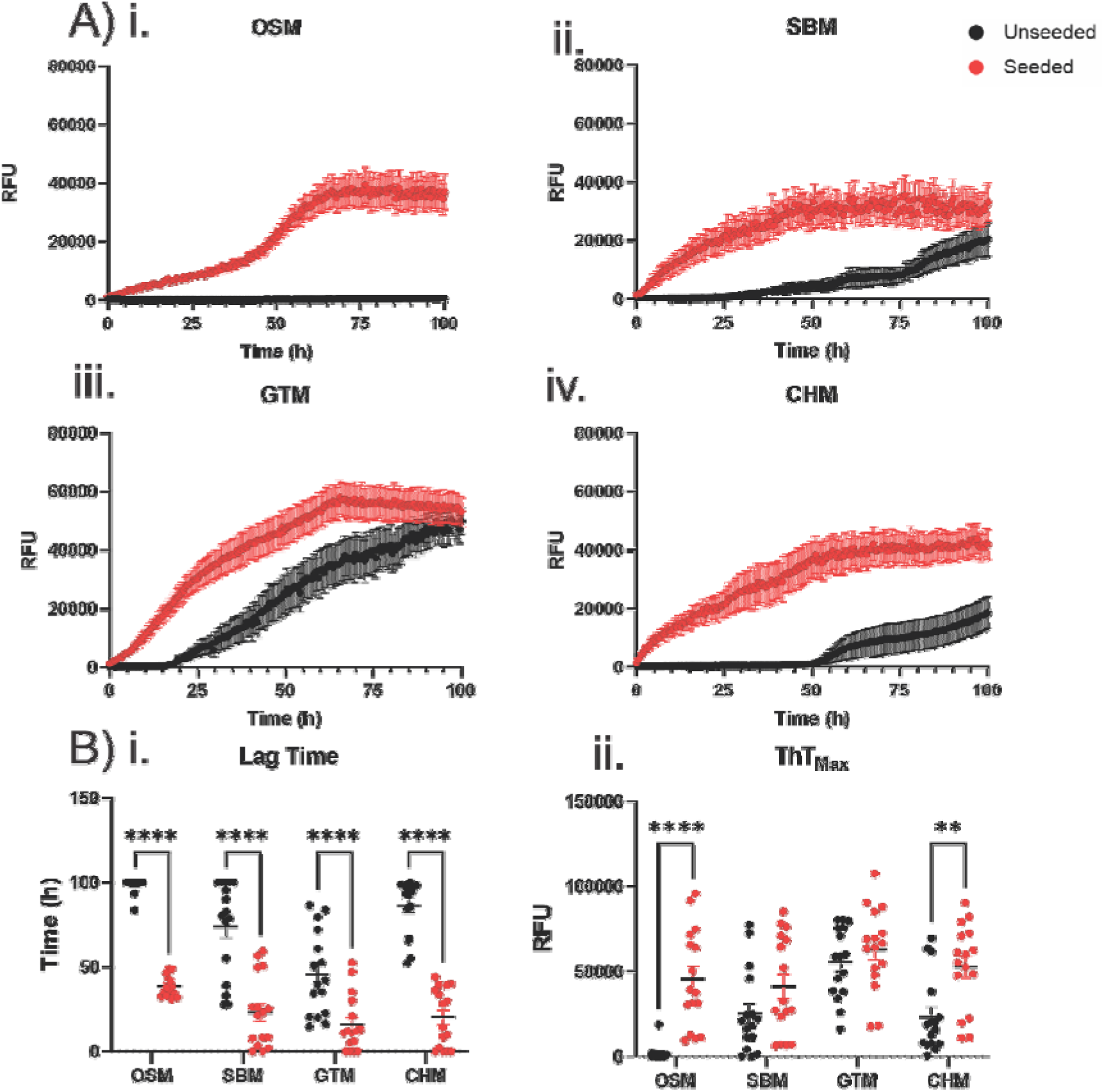
Comparison of Unseeded and PFF Seeded SAA Reactions. **A)** Overlay of the Osmotic Shock Monomer (OSM) (i), Sonicated and Boiled Monomer (SBM) (ii), GST-(iv) combined reactions from the unseeded and seeded experiments. **B)** Comparison of the lag phase (i) and *ThT_Max_* (ii) between the unseeded and seeded reactions. *P* values for Lag Time (i) were calculated with multiple Mann-Whitney test with a Holm-Šidák correction for multiple comparisons. *P* values for *ThT_Max_* (ii) were calculated using an Ordinary two-way ANOVA test with Šidák multiple comparisons test.

To determine the substrate that would be the best for SAA, the average lag time for each of the four independent experiments for the unseeded and seeded reactions was calculated for each substrate. A Kruskal-Wallis test was used to determine if the means of the four independent experiments differed significantly. If the means of the four independent experiments were significantly different, that would indicate significant inter-plate variability (i.e. the lag time was significantly different between plates). For the SBM and GTM substrates both the unseeded and seeded reactions showed significant differences in the means indicating inter-plate variability (Fig. 5A). In comparison, the OSM substrate showed no significant differences between the average lag time of the unseeded reactions, and no significant differences between individual plates in the seeded reactions. It should be noted that, in the OSM seeded reactions, Dunn’s multiple comparison test showed no significant differences between independent experiments (Fig. 5A). The CHM substrate showed no significant differences in either unseeded or seeded reactions (Fig. 5A). To further compare the substrates, the coefficient of variation (CV) was compared between the unseeded and seeded reactions for all four substrates. The CV measures the amount of variation in the lag times for each of the independent experiments. The greater the CV the greater the variation within an individual experiment (i.e. greater intra-plate variability in lag time time). A two-way ANOVA indicated that there were significant differences in the CV between the seeded and unseeded reactions, with the seeded reactions being associated with greater intra-plate variability. Tukey’s post hoc testing indicated that the CV of CHM unseeded and seeded reactions were significantly different, with the seeding reactions having higher CV (more intra-plate variability) compared the unseeded reactions (Fig. 5B). Furthermore, the seeded CHM reactions had greater CV than the seeded OSM reactions (Fig. 5B). This indicates that the CHM substrate, while not being associated with inter-plate variability, has significant intra-plate variability. In summary, limited inter- and intra-plate variability was observed with the OSM substrate while maintaining a significant difference between the lag times for seeded and unseeded reactions, a combination of properties not observed for other substrates.

**Figure 5:**
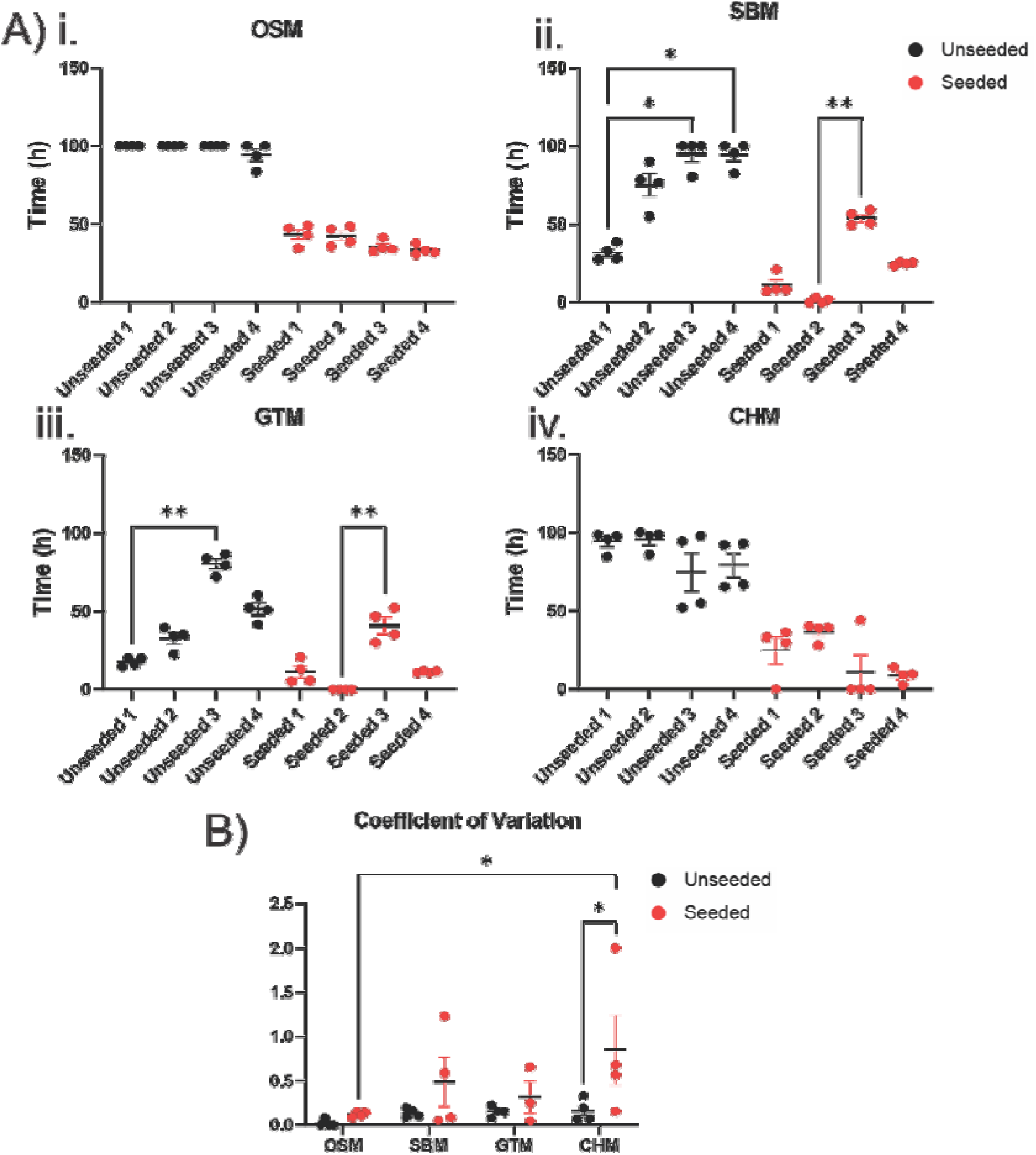
Substrate Variability for Unseeded and PFF Seeded SAA Reactions. **A)** The mean reaction time for each of the four independent quadruplicates for the Osmotic Shock Monomer (OSM) (i), Sonicated and Boiled Monomer (SBM) (ii), GST-Tagged Monomer (GTM) (iii), and Commercially Available His-Tagged Monomer (CHM) (iv) substrates. Kruskal–Wallis test followed by Dunn’s multiple comparisons test was used to calculate P values. **B)** The coefficient of variation was calculated for each of the independent experiments for both unseeded and seeded reactions for each substrate to determine the amount of inter-plate variability. *P* values were calculated via an Ordinary Two-Way ANOVA test followed by Tukey’s Multiple Comparison test.

### Specificity of Seeding in Brain Homogenate SAA is Dependent on Substrate

As SAA has been increasingly used for diagnosing synucleinopathies, we tested the substrates in reactions seeded with brain homogenate (BH), a biologically relevant seed source, at one of the laboratories. BH from the cerebellum of an MSA case was used as a positive (αSyn aggregate-containing) sample, while BH from a wild-type C57BL/6 mouse was used as a negative sample. In order to investigate the detection window, i.e. the difference between a positive and a negative BH sample, we calculated the lag time and *ThT_Max_* values of the SAA reaction. The difference in C57 and MSA lag time for GTM substrate was 0.85 h (C57: 15.78 h; MSA: 14.93 h) while SBM substrate was 2.83 h (C57: 19.74 h; MSA: 16.92 h) precluding their usage in SAA as the difference between positive and negative samples was not significant (Fig. 6A, B). The lag time for CHM substrate (Difference: 23.37 h; C57: 59.03 h; MSA: 35.66 h) and OSM substrate (Difference: 33.62 h; C57: 67.03 h; MSA: 33.41 h) showed significant differences (*p*<0.0001 for both substrates) between the MSA and C57 samples (Fig. 6B). To further highlight the OSM as the ideal substrate for SAA, only the OSM showed a significant difference (*p*<0.05) in *ThT_Max_* (Difference: 28,857; MSA: 34,809; C57: 5,953), however, this effect was likely primarily driven by incomplete aggregation in the C57 samples (Fig. 6B). These experiments were only conducted until 72h, unlike the 100h in the PFF SAA, as the reaction conditions differ. The OSM provides the most suitable substrate for SAA as all measured ThT kinetic parameters are different between positive and negative cases.

**Figure 6:**
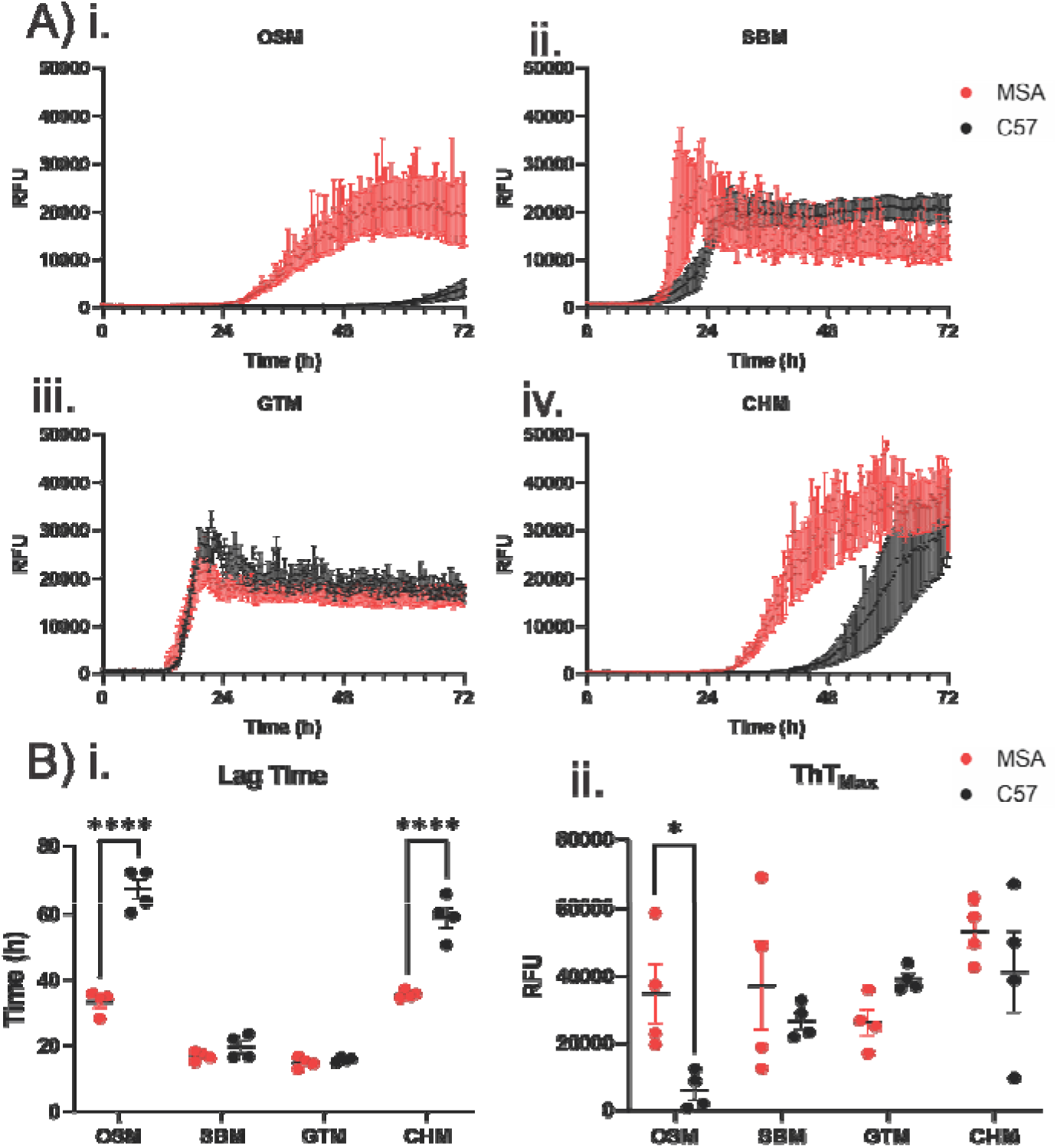
αSyn Substrate Performance in Brain Homogenate-Seeded SAA Reactions. **A)** Osmotic Shock Monomer (OSM) (i), Sonicated and Boiled Monomer (SBM) (ii), GST-Tagged Monomer (GTM) (iii), and Commercially Available His-Tagged Monomer (CHM) (iv) were seeded with 5µg of brain homogenate. **B)** Comparison of the lag phase (i) and *ThT_Max_* (ii) of all the seeded reactions. *P* values were calculated using a two-way ANOVA test with Šidák multiple comparisons test.

### OSM’s Resistance to Spontaneous Aggregation is Consistent Across Buffer Conditions

It has been widely reported that buffer conditions can affect the seeding response in αSyn SAA [21]. Therefore, we sought to investigate whether OSM’s low *de* novo aggregation potential was a consistent property of the purified protein or due to the buffer conditions and experimental setup. If OSM was resistant to *de* novo seeding, then unseeded reactions should follow similar trends regardless of buffer choice. For this purpose, we performed seeded and unseeded SAA reactions using a variety of different buffer conditions using OSM as the substrate. In fact, we observed OSM exhibited limited changes in unseeded reactions across the eight buffer conditions tested, suggesting low *de novo* aggregation propensity is a consistent property of the OSM substrate (Supplemental Fig. 2A, B). However, seeded reactions showed significant variation between buffers, suggesting seeded reactions are highly susceptible to changes in buffer conditions, which is aligned with existing literature. Specifically, altering the buffer had a notable impact on well-to-well variability.

### Characterization of αSyn Substrate Preparations

To explore potential explanations for their variable performance in the SAA experiments, the recombinant αSyn substrates were analyzed by SDS-PAGE followed by Coomassie blue staining. The OSM, GTM and SBM substrates all showed a single band at ∼16 kDa corresponding to full-length IZSyn (Fig. 7A). The CHM substrate showed a single band at ∼17 kDa, with the additional molecular weight likely corresponding to the N-terminal His-tag (Fig. 7A). While all substrates were normalized to 0.5 mg/mL prior to running on the gel, the CHM bound significantly more Coomassie blue, which we hypothesize may represent stronger binding of negatively-charged Coomassie blue to the positively-charged His residues within the His-tag. As the substrates all appeared comparably free from protein contamination, we next assessed the circular dichroism (CD) spectra of the proteins to look for differences in protein folding that could explain the results. The CD spectra of the GTM, SBM, and CHM substrates were consistent with random coil structure, with local minima around 200 nm (Fig. 7B). However, the OSM substrate showed a distinct right shift and broadening of the peak which may suggest some level of α-helical structure (Fig. 7B). Three readings of each OSM GTM, SBM and CHM, substrates were compared and OSM had a significantly (*p*<0.05) higher wavelength of the minima compared to all other substrates (GTM: 199.5; CHM: 196.4; SBM: 200.0; OSM: 204.9) (Fig. 7C).

**Figure 7:**
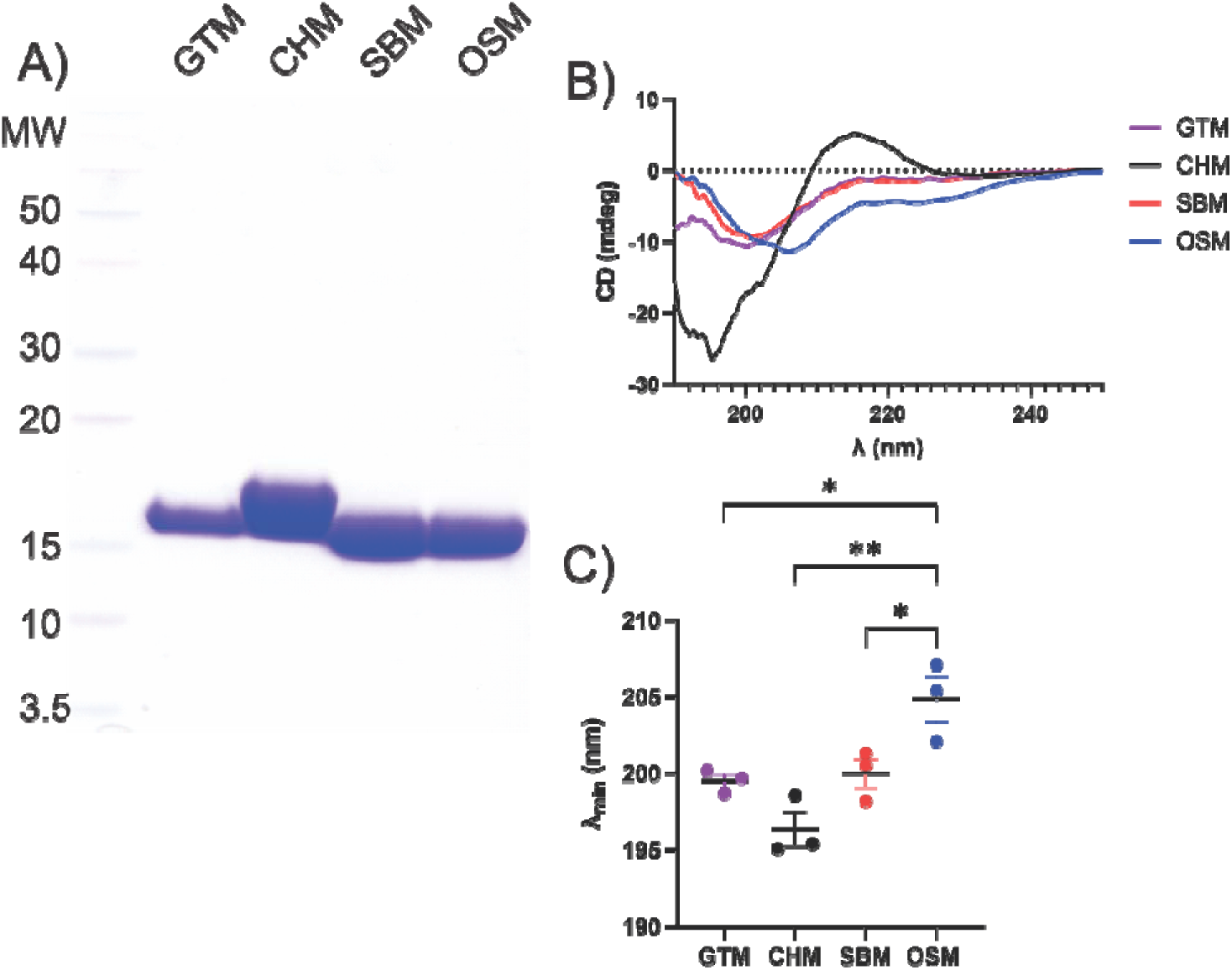
Comparison of αSyn Substrate Properties. **A)** All 4 substrates (15 µg per lane) were run on SDS-PAGE followed by Coomassie blue staining to assess potential protein contamination. **B)** The CD spectra of all four αSyn substrates were measured to compare their structural properties. The SBM, OSM, and GTM were averaged across three separate batches, whereas the CHM was averaged across two separate batches. C) Comparison of the wavelength minima in the CD spectra for OSM, SBM, GTM and CHM. *P* values were calculated using an Ordinary One-way ANOVA with Tukey’s multiple comparisons test.

### Protein Mass Spectrometry and Analytical Ultracentrifugation of Monomeric Substrates

We also analyzed purified monomer OSM, SBM, GTM and CHM using intact protein mass spectrometry and analytical ultracentrifugation to further understand potential differences that may explain differences in seeding propensity in SAA. Mass spectrometry analysis showed high consistency in expected mass values for different batches across the different purification methods (Fig. 8A). For each sample set, purity was consistent across both batches and the deconvoluted masses correlated well the expected amino acid sequences (< 1 Dalton tolerance): OSM, (theoretical: 14,460.2); SBM, (theoretical: 14,460.2); CHM, (theoretical: 15,584.4 – Met1 = 15,453.2); GTM, (theoretical: 14,871.6). Notably, the sonicated ‘SBM’ batches contained an extra peak at 14,704.7 (+245.3 mass shift) which was not present in the OSM batches. Common to bacterial expression, loss of the initiator methionine (-131.2 mass shift) and a minor secondary peak at 15,630.6 (expected +178.2 gluconoylation of His-tag) were detected for the CHM batches.

**Figure 8:**
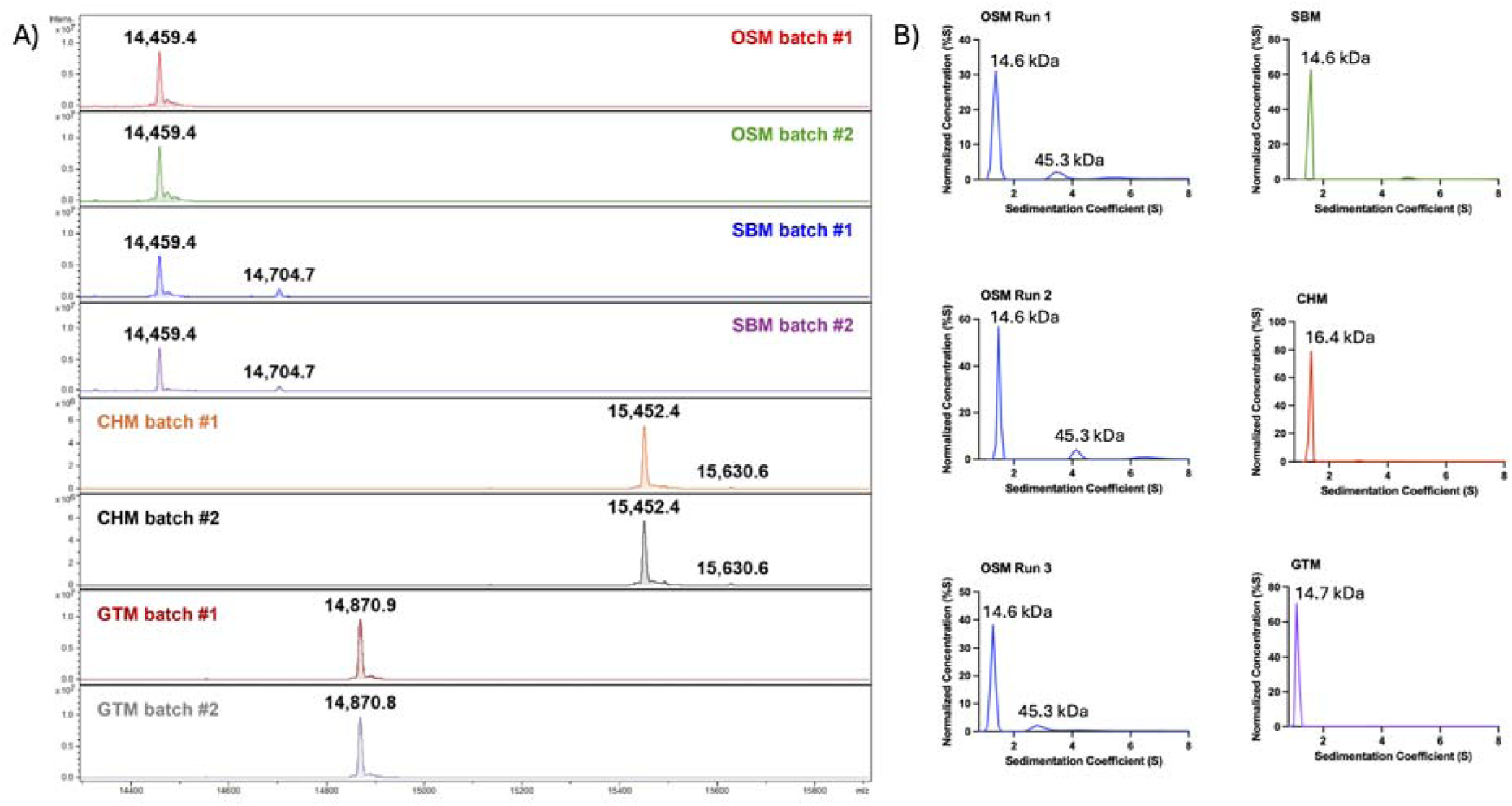
Mass Spectrometry and Analytical Ultracentrifugation Analysis of Substrates. **A)** Two batches each of OSM, SBM, CHM and GTM were analyzed using LC-MS to look for the presence of post-translational modifications. **B)** Sedimentation velocity analysis of two batches of OSM (over 3 runs) and one run each of SBM, CHM and GTM substrates were analyzed using absorbance optics. Concentration values were normalized using the sum of all values to enable comparisons of samples across different runs. Sedimentation coefficient values were plotted from 0.5 to 8.0 S and calculated molecular weights are shown.

In analytical ultracentrifugation experiments, all monomers showed a main sedimentation peak at the approximate molecular weight expected for each protein sequence (OSM: 14.4 kDa; GTM: 14.7 kDa; SBM: 14.6 kDa; CHM:16.4 kDa) (Fig. 8B). However, on three separate runs using two separate batches of OSM, a minor peak (∼15% of sample) corresponding to higher molecular weight (∼45 kDa) species was noted. This species was not observed in any of the other monomer preparations. This may reflect a tendency for OSM to form multimeric structures that are more resistant to aggregation in SAA leading to a lower *de novo* seeding propensity. However, deeper and more systemic analyses are required to understand the relevance and structural composition of these conformations.

## Discussion

As the use of αSyn SAAs continues to increase, so too does the need to standardize conditions between labs and to understand the potential impact different protocols can have on the results. In fact, researchers have recently been emphasizing the need for standardization of protocols since SAAs notoriously suffer from high variability [28–30, 36, 48]. This variability can have significant implications on the validity of the assay results for different biospecimens, especially as SAAs begin to move towards adoption in the clinical realm. To this end, we tested four recombinant αSyn substrate preparation methods for their viability to be used in SAA and compared the results obtained across two laboratories. The OSM substrate always displayed the most consistent results across all ThT kinetic parameters tested in unseeded and seeded reactions. Interestingly, limited batch-to-batch variability in OSM substrate has been reported by other groups when using cerebrospinal fluid as the seed for SAA [28]. These properties make OSM the ideal substrate for SAA as it reduces the rates of false positive reactions, while still allowing high sensitivity in seeded reactions and a high degree of intra- and inter-laboratory reliability.

As a substrate, GTM was not suited for SAA as it had the fastest *de novo* aggregation rate of all substrates tested, leading to false positive reactions that would make seeded reactions difficult to interpret. Moreover, GTM was found to be the fastest aggregating substrate in both laboratories, showing a high degree of intra- and inter-laboratory replicability. Interestingly, the CHM and SBM substrates both showed significant inter-laboratory variability. In one laboratory, the CHM substrate aggregated extremely quickly making it unsuitable for SAA, whereas the SBM substrate showed a low aggregation propensity comparable to OSM. However, the other laboratory found the exact opposite, with the SBM being a fast-aggregating substrate compared to the slow-aggregating CHM. It has been noted in the literature that some preparations of His-tagged IZSyn monomer can produce “slow” or “fast” kinetics batches of substrate [22]. This further supports the idea that batch-to-batch variability in certain preparations of substrate can be a significant concern in SAA [28, 30]. This is particularly important to consider when testing biological samples, as this may increase the rates of false positives or negatives if an assay was optimized using a “fast” kinetic batch but later run with a “slow” kinetic batch, or vice versa. Similarly, the choice of buffer conditions and cycle settings are pivotal in the final seeding kinetics achieved. Changing the buffers while using OSM had a significant impact on lag phase, well-to-well variability, and *ThT_Max_* value in seeded reactions. Therefore, it may be prudent to characterize the kinetics of each batch of substrate prior to further use.

In an attempt to understand why the various recombinant αSyn substrates behaved differently in the SAA, we explored multiple possibilities. As the behavior of SBM and OSM substrates, which both consist of full-length untagged wildtype IZSyn, were different, we think it is unlikely that simple sequence difference could explain seeding differences observed. Furthermore, as the four substrates each displayed a single band following SDS-PAGE, we do not believe that protein contamination is a likely explanation.

A unique biophysical property of the OSM substrate was that the peak of the CD spectra was right shifted and broadened compared to the other substrates. We speculate that this could indicate some proportion of α-helical content in the OSM not present in the other substrates. This shift in the CD spectra of OSM compared to the other substrates mirrors the CD spectra changes noted in tetrameric compared to monomeric αSyn by other groups [49, 50]. Additionally, using analytical ultracentrifugation, we noted that OSM preparations contain a minor species with a higher molecular weight, raising the possibility that OSM αSyn monomers may exist in equilibrium with higher-order structures. It has been postulated that α-helical tetrameric αSyn species are aggregation-resistant whereas monomeric unstructured αSyn is aggregation-prone [49, 50]. However, this resistance to aggregation only applied to *de novo* aggregation, as tetrameric αSyn could still be seeded as effectively as monomeric αSyn [49, 50]. Tetrameric αSyn can undergo a temperature-dependent irreversible-dissociation, which may explain why the SBM is a poor substrate for SAA, as the boiling step in the purification would likely destroy tetrameric αSyn species [49, 50]. The presence of either an N-terminal poly-histidine tag or the residual N-terminal sequence ‘GPLGS’ following cleavage of the GST-tag in CHM and GTM, respectively, could alter either the ability of αSyn to form aggregation-resistant tetramers or their equilibrium with monomeric αSyn, which could hinder their use in SAA [49, 50]. Finally, previous work involving purification of tetrameric αSyn used non-denaturing conditions, similar to the OSM protocol, which could imply that tetrameric αSyn species are preserved in OSM [49, 50].

Another potential explanation for the differential substrate behaviors may be the oxidation state of IZSyn. It has previously been reported that αSyn can become oxidized on its four methionine residues, resulting in the formation of methionine sulfoxides [51, 52]. Oxidized αSyn has a lower *de novo* aggregation propensity compared to unoxidized αSyn [52]. Osmotic shock is often called periplasmic lysis, as it selectively harvests proteins from the periplasmic space opposed to the cytosol [37, 53]. However, the reported changes in CD spectra of oxidized αSyn are not identical to what we found for the OSM substrate [52]. In addition, mass spectrometry analysis did not indicate the presence of oxidation in two separate purifications of OSM. A further challenge to this hypothesis is that previous papers reporting “fast” and “slow” kinetic recombinant αSyn batches use a modified osmotic shock procedure. However, that study used His-tagged IZSyn, which may be a confounding factor given that we observed variable results with our CHM substrate [22]. This is further supported as the CHM used in this study was also produced via a modified osmotic shock procedure and similarly showed two very different kinetic profiles between the batches tested in each lab. It is also possible the explanation for why each different substrate did not perform as well may not be a common factor and instead could be due to unique factors specific to each purification method.

## Conclusion

As the field continues towards the use of SAA for the diagnosis and prognosis of synucleinopathies, care must be taken in the optimization and design of these assays to ensure intra- and inter-laboratory reproducibility. This is especially important as these technologies begin their transition into the commercial and clinical realms. In this work, we demonstrate that the method used to purify the αSyn substrate can dramatically impact its SAA performance. We therefore strongly encourage the use of OSM as the universal substrate for αSyn SAA.

## List of abbreviations

αSyn: α-Synuclein
PD: Parkinson’s disease
MSA: Multiple system atrophy
DLB: Dementia with Lewy bodies
SAAs: Seed amplification assays
ThT: Thioflavin T
AD: Alzheimer’s disease
GTM: GST-tagged IZSyn monomer
SBM: Sonication and boiling αSyn monomer
OSM: Osmotic shock αSyn monomer
CHM: Commercial His-tagged αSyn monomer
BH: Brain homogenate
CV: Coefficient of variation
CD: Circular dichroism

## Declarations

### Ethics approval and consent to participate

Human α-synucleinopathy samples were provided by the Massachusetts Alzheimer’s Disease Research Center. Informed consent was obtained either by the patient prior to death or by family member prior to or at the time of death. The use of human tissue was in accordance with guidelines provided by the University of Toronto, and all relevant ethical regulations were followed.

### Consent for publication

Not applicable.

### Availability of data and material

All data generated or analysed during this study are included in this published article [and its supplementary information files]. Raw SAA kinetic curves can be made available upon reasonable request from the corresponding authors.

### Authors’ contributions

ZAMA and NRGS contributed equally to the collection, analysis, and interpretation of all results included within then manuscript. SN aided in the performance and optimization of OSM purification methodology. GTM purification and PFF generation were performed by WL. Electron microscopy characterization and dynamic light scattering analyses were performed by WL and IS, respectively. MI and BTH generously provided the brain homogenate samples used in this study. JFT led the mass-spectrometry and ultracentrifugation result analyses and interpretation for all monomer substrates. GTM and PFF batches were kindly provided by TMD. ZAMA, NRGS, JCW and EAF were responsible for the study conception and design. First draft was written by ZAMA and NRGS. All authors read and approved the final manuscript.

## Acknowledgements

The authors thank Mark A. Hancock (McGill SPR-MS Facility) for assistance with the intact protein LC-MS analyses, for which infrastructure was gratefully provided by the Canada Foundation for Innovation (CFI). The authors thank Kim Munro, staff scientist at the centre de recherche en biologie structurale, for conducting the analytical ultracentrifugation analysis, which was funded by Fonds de Recherche du Québec (Health Sector) Research Centres Grant #288558.

