## Supplemental for "α-Synuclein purification significantly impacts seed amplification assay performance and consistency"

### Supplementary Material

#### Supplemental Methods

##### Characterization of PFFs using transmission electron microscopy

For EM analysis, PFFs were diluted in dH2O to a final concentration of 20 µM, with 5 µl of the PFF mixture loaded onto a carbon-coated copper grid and left to sit for 2 minutes. Next, it is fixed with 5 µl of 4% paraformaldehyde (PFA) for 1 minute, before the PFA is removed and the grid is washed with dH2O for 1 minute. The washes with dH2O are repeated for an additional 3 washes. Finally, the PFFs on the grid are stained with 2% uranyl acetate for 1 minute, before being removed and air dried for 30 minutes. Images were acquired using the Tecnai Spirit TEM after negative staining of PFF samples fixed on the carbon-coated copper grids as described previously.[54]

##### Characterization of PFFs using dynamic light scattering

PFFs of α-syn were generated as previously described.[40,55,56] The generated fibrils were then sonicated using Bioruptor Pico sonicator (Diagenode) for at least 40 cycles of 30 sec on/30 sec off program to achieve ≤100 nm sizes. Sizes and morphology of PFFs were monitored by electron microscopy and dynamic light scattering (DLS). For DLS analysis 0.5-1mg/mL α-syn PFFs solution was prepared by dilution in filtered (at 0.1 micron) PBS and centrifugation at 13000 rpm for 5 min. The supernatant was transferred to DLS cuvette and analyzed using the Zetasizer Nano S system (Malvern Panalytical).

### Supplemental Figures

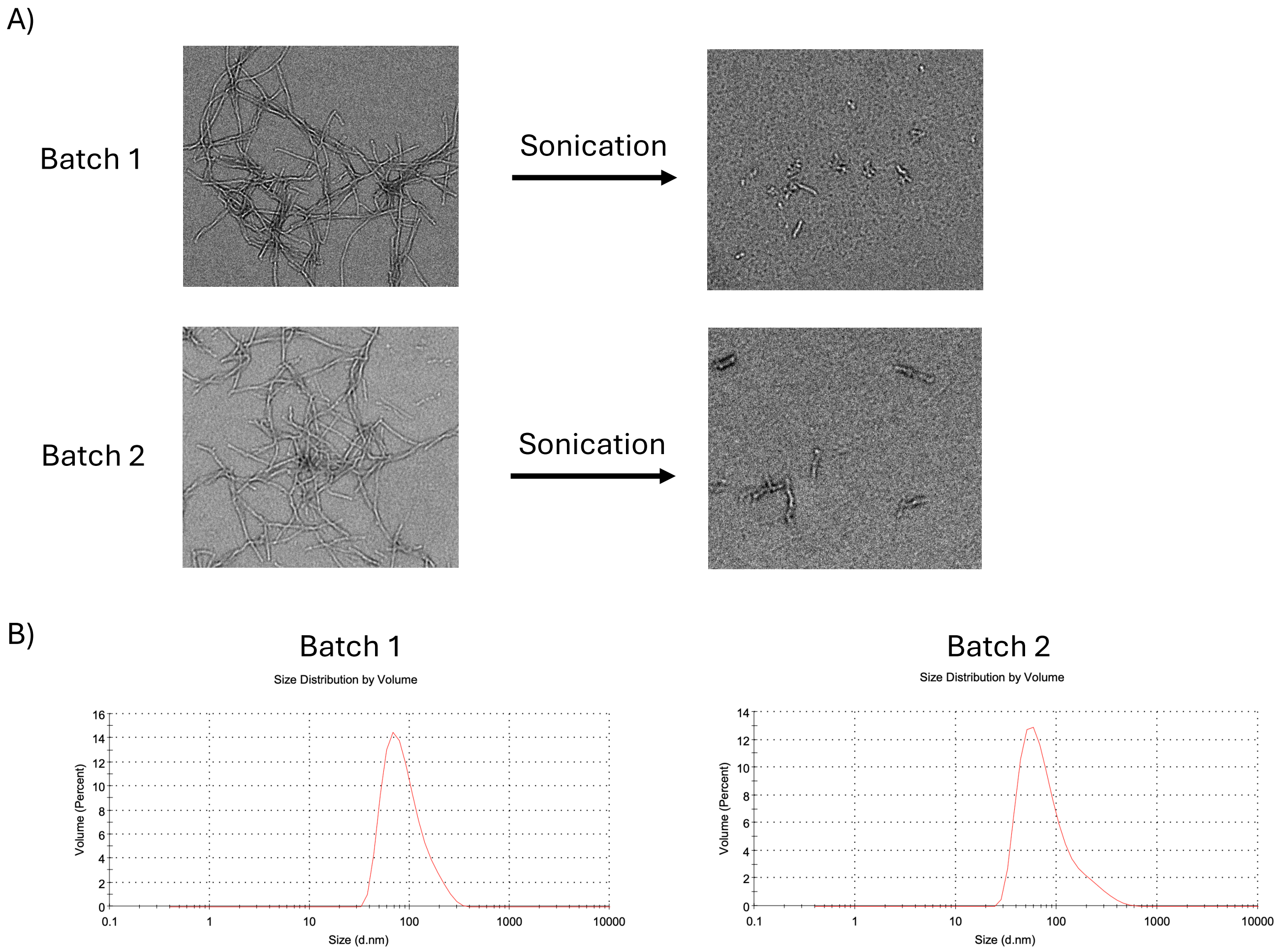


**Supplemental Figure 1: Structural Characterization of PFFs. A)** Transmission electron microscopy (TEM) performed on two separate PFF batches before and after sonication. **B)** To ensure homogeneity after sonication, dynamic light scattering (DLS) analysis was performed on each batch. Hydrodynamic diameter measurements were 103.5 nm and 103.2 nm for batch 1 and batch 2, respectively.

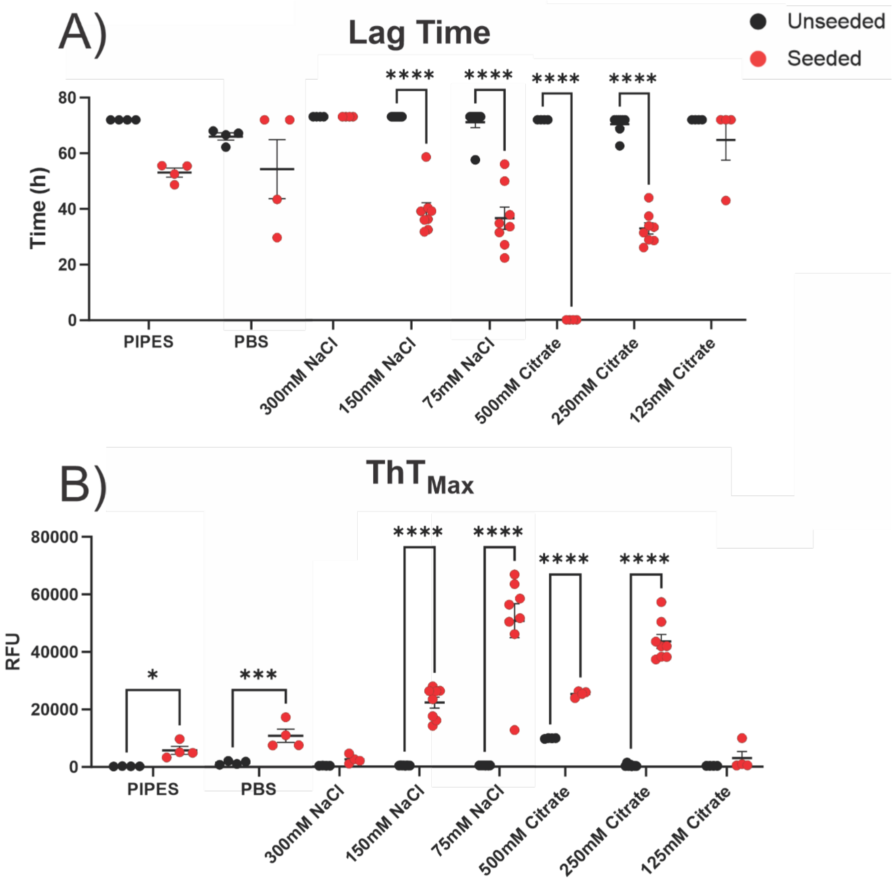


**Supplemental Figure 2: Different Buffers Impact Seeded but Not Unseeded SAA Reactions Using OSM Substrate.** Seeded and unseeded reactions using eight different buffers' lag times **(A)** and $ThT_{Max}$ **(B)** were calculated. Experiments were run for a maximum of 72h. If aggregation did not occur in this timeframe a lag time of 72h was assigned to the well. *P* values for Lag Time were calculated with a Mann-Whitney test with a Two-Stage Linear Step-up Procedure and $ThT_{Max}$ were calculated with Ordinary Two-Way ANOVA test followed by Tukey's Multiple Comparison test.
